# Deep learning using structural MRI massively improves prediction accuracy of body mass index

**DOI:** 10.1101/2025.04.11.648433

**Authors:** Alysha Cooper, Mahmoud Elsayed, Max Owens, James MacKillop

**Affiliations:** Peter Boris Centre for Addictions Research, St. Joseph’s Healthcare Hamilton, Hamilton, ON, Canada; Department of Psychiatry and Behavioural Neurosciences, McMaster University, Hamilton, ON, Canada

**Keywords:** Deep learning, convolutional neural networks, obesity, brain morphometry, explainable AI

## Abstract

Obesity is a major public health problem globally and there is considerable interest in the neural mechanisms in food overconsumption. Artificial intelligence (AI), particularly machine learning, has shown promise in characterizing links between brain morphometry and obesity. In 1106 adults, compared to other forms of machine learning, deep learning using 3D convolutional neural networks (3D-CNN) dramatically improves prediction of body mass index (BMI). The 3D-CNN model robustly predicted BMI (*R*^2^ =.325), outperforming random forest, elastic net, and tabnet models (*R*^2^s<.07) in a ‘lockbox’ sample. Explainable AI analyses revealed the specific brain regions implicated and these regions were moderately associated with delay discounting, fluid cognition, gait speed, dexterity, and alcohol use. Collectively, these findings reveal the value of deep learning for understanding of the neural basis and motivational processes in the neurobiology of obesity.

## 1 Introduction

Over recent years, machine learning has grown in popularity for predicting health outcomes to aid in clinical decision making (Chattopadhyay & Maitra, 2022; Cho et al., 2021; Ganggayah et al., 2019; Iwase et al., 2022; Loutati et al., 2024). Unlike traditional statistical models, machine learning methods often avoid relying on distributional assumptions, linearity assumptions, and/or multicollinearity assumptions. Perhaps most important, machine learning methods can incorporate vast numbers of predictors in a dataset (Rajula et al., 2020). Despite these potential advantages to machine learning over more traditional statistical methods, results are mixed in terms of the comparison in prediction performance between the two frameworks in applications to health. For instance, some studies find comparable performance between machine learning methods and traditional statistical models in the prediction of health outcomes (Desai et al., 2020; Li et al., 2020) while others find notable improvement when using machine learning methods instead of statistical models (Ganapathy et al., 2022).

Tomographic imaging methods such as magnetic resonance imaging (MRI) are commonly used for diagnostic or disease clarification purposes. While more common machine learning methods such as random forest and regularized regression have been used to predict health-related outcomes using atlas-based features based on medical images (Jollans et al., 2019), they lack the ability to engineer (i.e., extract) the features themselves from the raw data. On the other hand, deep learning provides methods that are capable of learning data representations directly from images. Specifically, the most commonly used method for computer vision tasks is the convolutional neural network (CNN), introduced by LeCun et al. (1989), which are specialized neural networks with layers that perform convolution operations to create feature maps. In each convolution layer, the CNN will create a variety of filters (i.e., small matrices) which are used to detect the presence and strength of features or patterns in an image. In the first few layers of the CNN, these filters will detect higher-level features such as edges and textures, but the features become increasingly complex and uninterpretable as the layers progress. Unlike most standard machine learning methods, CNNs can capture spatial dependencies and demonstrate the ability to recognize objects regardless of their location in an image (Albawi et al., 2017) making them useful in imaging diagnostics.

Obesity, an endocrine disorder characterized by excess adipose tissue, continues to grow worldwide with a four-fold increase in prevalence among adolescents and a two-fold increase in prevalence among adults from 1990 to 2022 (World Health Organization, n.d.). These noticeable increases in prevalence are of concern due to obesity being a risk for mortality (Organization & others, 2009), cardiovascular disease (Ortega et al., 2016), and other comorbidities including sleep apnea, stroke, and impaired fertility to name a few (Wilborn et al., 2005). Importantly, although adipose tissue is part of the endocrine system, a fundamental cause of obesity is excess energy imbalance (i.e., higher caloric consumption than expenditure) and food motivation is substantially regulated by the brain.

Several previous studies have demonstrated associations between the brain structure and BMI via machine learning approaches. For example, in contrast to early work that did not show advantages of machine learning (Syan et al., 2019), Xu, Owens, and MacKillop (Xu, Owens, et al., 2023) demonstrated superior performance in the prediction of BMI for participants when using an elastic net model with structural MRI data as opposed to a model with only demographic features. These findings suggested an association between BMI and regions of interest in the brain including the central executive network, default mode network, and salience network. Further analyses showed that the effect of the predicted BMI (i.e., neuroanatomical profile of BMI) on actual BMI was partially mediated by impulsive delay discounting (overvaluation of immediate rewards) and overall cognitive ability, converging with the larger behavioral literature linking obesity with delay discounting (Amlung et al., 2016; Tang et al., 2019), reward sensitivity (Sutton et al., 2022), and aspects of mental health (Avila et al., 2015), such as ADHD (Martins-Silva et al., 2021). Similarly, in children, multiple studies using machine learning found cortical surface area, cortical thickness, and gray matter volume to be predictive of BMI (Adise et al., 2021; Wang et al., 2023). However, a limitation to these approaches is that they relied on pre-engineered atlas-based features measuring volume, thickness, area, and surface area of the brain and did not use methods capable of learning representations directly from the structural MRI images. To date, only two studies have used deep learning to extract high-dimensional features from MRI images for prediction of obesity-related phenotypes (Guan et al., 2020; Vakli et al., 2020), one in children and another in adults. Both implicated a number of motivationally relevant brain regions and demonstrated the benefits of using deep learning for prediction of obesity outcomes. However, in both studies, it was not clear how sensitive the findings were to the specific train/test split given that neither study conducted cross-validation for model evaluation. Further, the 3D-CNN fit by Vakli et al. (2020) was not tested against other deep learning or machine learning models leaving its comparative performance unclear. Overall, these studies provide initial proof of concept that BMI can be predicted using deep learning analysis of structural MRI data, but several substantive open questions remain.

The ability to predict BMI from structural MRIs has important potential clinical applications. First, if BMI can accurately be predicted with structural MRIs alone, this would reflect an important relationship between the brain structure and BMI. Developing a better understanding of this relationship can help to determine potential biological causes of obesity. Using explainable AI methods to identify the important brain regions in prediction of BMI can add to the growing body of literature studying how structural changes in the brain are implicated in regulation of body weight. Allowing deep learning models to learn features directly from the raw imaging data is a promising approach for identifying potential neurobiological determinants of obesity, paving the way for targeted treatments. Moreover, a difference in brain-predicted BMI and actual BMI might be an important predictor of future health outcomes such as weight gain/loss or eating disorders. This application would be similar to the brain-age gap literature which looks at how the difference between brain-predicted age and actual age is an important predictor of neurodegenerative, neurological, and neuropsychiatric diseases (Franke & Gaser, 2019).

By applying a 3D-CNN to the prediction of BMI with structural MRIs from the Human Connectome Project, we: 1) demonstrate a strong association between the brain and BMI, 2) demonstrate the advantages of using of a 3D-CNN for feature extraction compared to other machine learning methods relying on pre-engineered atlas-based features, and 3) identify brain regions and psychological mechanisms implicated by the 3D-CNN model. To build upon previous research, the prediction performance of the 3D- CNN was compared to another deep learning method, TabNet (Arik & Pfister, 2021), in addition to elastic net regression (Zou & Hastie, 2005) and random forest regression (Ho, 1995). TabNet is a deep learning model optimized with gradient-descent, specifically designed for tabular data, hence the name. Unlike most deep learning models, TabNet offers enhanced interpretability through a sequential attention mechanism that selectively chooses features at each decision step (Arik & Pfister, 2021). Furthermore, sequential boosting was conducted in which the 3D-CNN features were further used as predictors in additional machine learning analyses to allow for a direct comparison of data learned directly from the images versus using atlas-based ROI features based on the images. In an important contrast to previous studies, nested cross-validation was employed to establish generalizability of model performance across different train/test splits.

## 2 Methods

### 2.1 The Data

Data from the S1200 data release of the Human Connectome Project (HCP) were used in all analyses. Data from HCP includes various imaging modalities along with behavioural and genetic data for 1,206 subjects. Measurements for participants took place at Washington University, St. Louis, United States, over a two-day period sometime between 2012 and 2015. All subjects were between the ages of 22 to 35 at the time of the study and did not have any known psychiatric disorder, substance abuse, neurological, or cardiovascular disease. For more details on the inclusion and exclusion criteria of HCP, we refer the reader to Van Essen et al. (Van Essen et al., 2012). The institutional review board of Washington University approved the HCP. HCP structural MRIs (sMRIs) are registered to the MNI-152 space. We followed the methods of Abrol et al. (Abrol et al., 2021) for pre-processing the sMRI data. Specifically, after segmenting the sMRIs into tissue probability maps for grey matter, white matter, and cerebral spinal fluid using VBM in SPM12, the grey matter maps were warped, modulated, and smoothed using a Gaussian Kernel with a full width half max of 10mm. With voxel size 1.5x1.5x1.5mm^3^, the preprocessed grey matter volume images had dimensions of 121x145x121.

BMI was used as the target variable (i.e., dependent variable) for all models fit. The processed sMRI brain scans were converted into *Numpy* (Harris et al., 2020) arrays of size 121x145x121 to be used as predictors in the 3D-CNN. Subjects were removed for 3D-CNN analysis if they failed MRI quality control, had missing sMRI scans or missing BMI measurements. The final analytical sample for the 3D- CNN comprised 1,106 subjects. For the EN, RF, and TabNet models, demographic data (i.e., gender, age, education, income) and atlas-based features based on the sMRIs (i.e., thickness and surface area of each cortical region, grey matter volume of subcortical regions, and total intracranial volume) were used as predictors. Of the 1,106 subjects used for training and evaluation of the 3D-CNN, *n* = 7 had missing demographic data and were removed from analysis for the three comparison models.

### 2.2 Measures

#### 2.2.1 BMI

BMI data was collected alongside other physical and health measurements as part of the non-medical imaging data collection of HCP. Participants’ self-reported heights and weights were used to calculate BMI as follows:

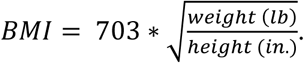

#### 2.2.2 Delay Discounting

Delay discounting was measured via two tasks asking participants to choose between a smaller, more immediate award versus a larger, delayed reward(Estle et al., 2006). Participants make the selection at 6 different delays (1 month, 6 months, 1 year, 3 years, 5 years, 10 years) for two delayed amounts ($200 and $40,000). For each delayed amount, the area under the curve discounting measure (AUC) is used to describe the steepness of delay discounting, with lower AUC reflecting greater delay discounting. The AUC has previously been shown to be a valid and reliable measure of delay discounting (Myerson et al., 2001).

#### 2.2.3 General Cognitive Ability

Two scores, the Fluid Cognition Composite score and the Crystallized Cognition Composite score, were derived via responses on the NIH Toolbox Cognition Battery which comprises neuropsychological tasks such as the Change Card Score Test and the Flanker Inhibitory Control and Attention Test (Weintraub et al., 2013). In the current study, we use the total Cognition Composite score (i.e., the sum of Fluid Cognition and Crystallized Cognition Composite scores) to measure general cognitive ability.

#### 2.2.4 Alcohol Use

The Semi-Structured Assessment for the Genetics of Alcoholism (SSAGA) (Bucholz et al., 1994) was administered during an over-the-phone assessment to collect self-reported information on substance use, mood, anxiety, eating disorders, and ADHD. Frequency of alcohol use in the past 12- months (4-7 days/week (1 if male, 2 if female), 3 days/week = 2, 2 days/week = 3, 1 day/week = 4, 1-3 days month = 5, 1-11 days/year = 6, never in past 12 months = 7) and whether or not the participant has met DSM4 criteria for DSM4 Alcohol Dependence at some point in their life (yes, no) were used to measure alcohol use in the current study.

#### 2.2.5 Mental Health

Depression and anxiety were assessed via the Achenbach Adult Self-Report (ASR) (Achenbach, 2009). Specifically, the ASR DSM Anxiety Problems raw score and the ASR DSM Depressive Problems raw score were used to measure anxiety and depression, respectively, in the current study. The ASR DSM Anxiety Problems scale and the ASR DSM Depressive Problems scale each comprise items that are consistent with the corresponding diagnostic category in the DSM (Achenbach et al., 2003).

#### 2.2.6 Motor Skills

In the current study, dexterity and gait speed were used as indicators of motor skills. Gait speed was measured using the NIH Toolbox 4-Meter Walk Gait Speed Test and dexterity was measured using the NIH Toolbox 9-Hole Pegboard Dexterity Test (Reuben et al., 2013). In the Gait Speed Test, participants were asked to walk 4 meters at their usual pace and gait speed was measured as time in seconds to walk the 4 meters. In the Dexterity Test, participants were asked to place and remove 9 plastic pegs from a plastic pegboard and their time was recorded.

#### 2.2.7 Atlas-Based Features

In the current study, the 255 atlas-based features included volume, thickness, and surface area measurements for both cortical and sub-cortical regions, based on the Conte69 atlas (Van Essen et al., 2012). The data were downloaded directly from HCP. For further details on the processing, we refer the reader to Glasser et al. (Glasser et al., 2013).

### 2.3 Predicting Body Mass Index

We used a 3D-CNN, random forest, elastic net, and TabNet to predict BMI. While the 3D-CNN was fit to the raw imaging data, the machine learning models and TabNet were fit to derived tabular atlas-based features. A summary of our methods is provided in Figure 1.

**Figure 1.**
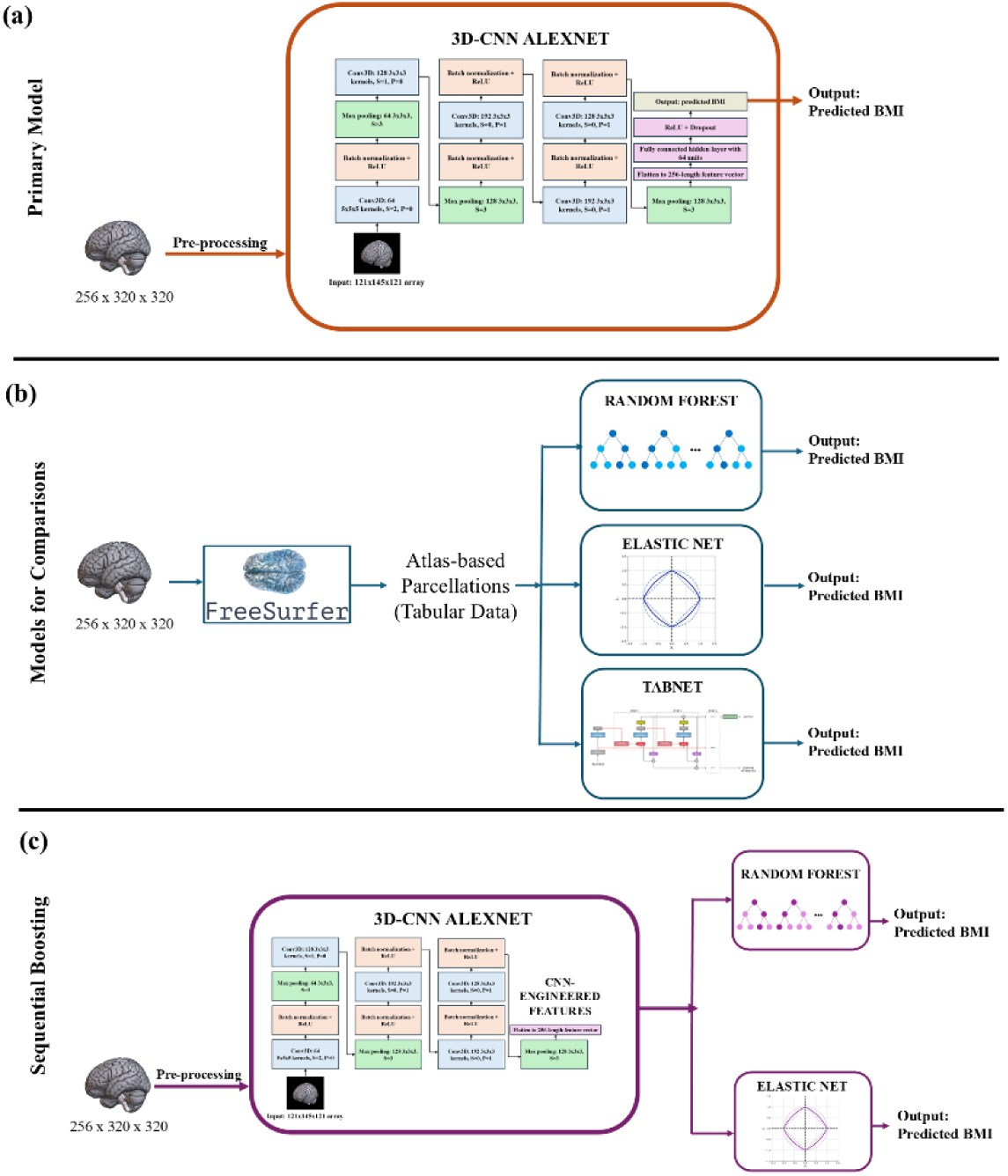
Schematic of models fit for prediction of BMI. Models fit included a 3D-CNN with the AlexNet architecture(Krizhevsky et al., 2012), TabNet(Arik C Pfister, 2021), elastic net(Zou C Hastie, 2005) and random forest regression(Ho, 1995).

#### 2.3.1 3D-CNN

Our primary model of interest was a 3D-CNN based on the AlexNet architecture (Krizhevsky et al., 2012) with five convolutional layers. Figure 2 presents the architecture of the network. The code was adapted from Abrol et al. (Abrol et al., 2021) and implemented with *PyTorch*(Paszke et al., 2019). The Adam optimizer (Kingma, 2014) was used to minimize the loss function, which was chosen to be the mean squared error (𝑀𝑆𝐸). Using this loss function allowed us to minimize the difference between predicted BMI and observed BMI. The learning rate of the Adam optimizer and the batch size were selected via hyperparameter tuning, which is described in more detail below. The learning rate could take on any value from 0.0001 to 0.05 while the batch size could be one of 8, 16, 32. Momentum was set to 0.9. A learning rate scheduler was employed to reduce the learning rate once there were no longer improvements in the loss function on the validation set. Three methods were used to reduce the risk of overfitting: 1) weight decay, 2) dropout, and 3) early stopping. Both the weight decay penalty and the dropout proportion were additionally selected through hyperparameter tuning with possible parameters ranging between (1e-4, 0.5) for weight decay and (0.3, 0.8) for dropout. Training was prematurely stopped if no improvement in reducing the loss function on the validation set was observed for 40 epochs. A maximum of 200 epochs was used. Model weights from the best performing epoch were used for model evaluation on the test set. Initial weight values of each CNN layer were sampled from the normal distribution with mean zero and standard deviation of 2/√𝐾, where 𝐾 is the total number of weights to be initialized in the layer. The weights of batch normalization layers were initialized with values of one while the biases were initialized with values of zero.

**Figure 2.**
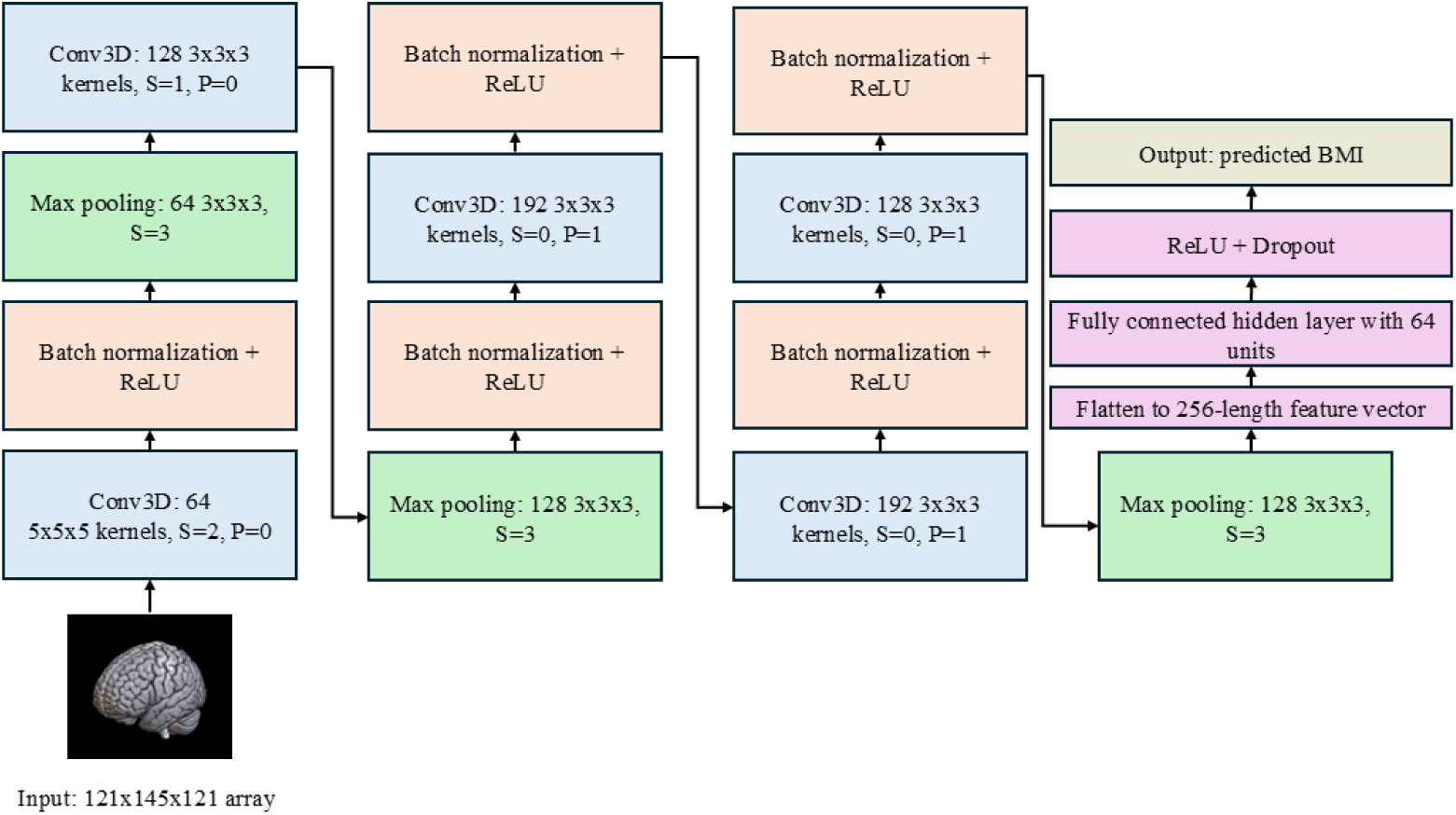
Schematic of architecture for 3D-CNN used to predict body mass index from predictor arrays of size 121x145x121. S denotes stride, P denotes padding.

#### 2.3.2 Models for Comparison

To assess whether the 3D-CNN model provided advantages in learning complex relationships beyond what can be learned from atlas-based features, we additionally fit the following models for comparison: TabNet (Arik & Pfister, 2021), random forest, and elastic net. Each of these three models are designed for tabular, structured data. The three models were fit three times: once with only demographic covariates (i.e., sex, age, years of education completed, household income, intra- cranial volume), once with only brain covariates (i.e., atlas-based), and once with both demographic and brain covariates. Fitting three separate models with different subsets of predictors allowed us to assess whether the atlas-based brain features contributed to model prediction beyond demographics alone. TabNet is a deep learning model comprising layers for attention transformers and feature transformers, allowing it to identify relevant features in addition to processing features into a more useful representation for prediction. The TabNet model was fit with the Pytorch implementation of TabNet. The tuned hyperparameters of TabNet included the learning rate, weight decay penalty, batch size, the number of features passed forward to the attention network (𝑛_𝑎_) and the number of features involved in the decision step (𝑛_𝑑_). As recommended by Arik and Pfister (Arik & Pfister, 2021), we tuned 𝑛_𝑎_ and 𝑛_𝑑_ together such that we set 𝑛_𝑎_ = 𝑛_𝑑_. Similar to the 3D-CNN setup, a learning rate scheduler was employed for TabNet as well, and the maximum number of epochs was set to 200 with early stopping after 40 epochs of no improvement. Model parameters from the best performing epoch were used for model evaluation on the test set. Both elastic net and random forest were fit with the *Scikit-learn* package (Pedregosa et al., 2011) in Python. For the random forest, we tuned the number of trees in the forest, the maximum depth of each tree, and the minimum number of samples to split. All other parameters of random forest were set to the default values in the *Scikit-learn* package. As for elastic net, we tuned the penalty parameter, 𝜆, to determine the appropriate balance between model fit and model complexity as well as the parameter, 𝛼, to determine the appropriate balance between the ridge penalty and the lasso penalty.

Finally, the standard machine learning models, random forest and elastic net, were fitted again using the engineered features from the trained CNN model for a sequential boosting approach. Using the CNN features to train random forest and elastic net allowed for us to evaluate whether any improved performance of the CNN was due to the combination of feature extraction as well as the feed forward neural network for prediction versus the improvement solely being explained by the feature extraction layers. If similar performance was seen across the random forest, elastic net, and CNN after using the CNN-extracted features, then it could be determined that the feature learning that CNN offers is what makes CNNs a powerful tool to better understand the relationship between the brain and BMI.

### 2.4 Model Fitting and Evaluation

The dataset was split into a training set comprising roughly 80% of the data and a testing set comprising the remaining 20%. Models were not evaluated on the test set until hyperparameter tuning was complete and the overall analysis pipeline was finalized and, therefore, we refer to this test set as the “lockbox set”. The train/test split was conducted on the Family ID variable to ensure that those within the same family would not be used to train models for prediction on others within the family given sibling oversampling in the HCP. On the training set, a 5-fold cross-validation (CV) procedure with three repeats was employed for evaluation of model fits to see how performance varied over different data splits. The mean evaluation metrics along with their standard deviation across the fifteen total folds were reported. For hyperparameter tuning, a nested 3-fold CV procedure was performed. For nested CV, each training CV fold was further divided into three folds to evaluate different combinations of hyperparameters. The hyperparameter combination which maximized the mean 𝑀𝑆𝐸 across the three nested folds was selected. Combinations of hyperparameters to evaluate were randomly sampled via random search. Finally, the best model across the fifteen outer folds was used for prediction on the lockbox set to further evaluate the model performance. To ensure fair comparison across models, the same data splits were used for the 3D- CNN, random forest, elastic net, and TabNet.

### 2.5 Explainable Artificial Intelligence

#### 2.5.1 Gradient Regression Activation Mapping

We used an alternative form of gradient class activation mapping (GRAD-CAM) (Selvaraju et al., 2017) applicable to regression tasks to better understand which regions of the structural MRIs were relevant to the 3D-CNN predictions. From hereon, this method will be referred to as GRAD regression activation mapping (GRAD-RAM). To identify image regions important to the prediction, GRAD-RAM uses the gradients flowing into the final convolutional layer during backpropagation. We computed the gradient of the predicted BMI with respect to each feature map, 𝐴^𝑚^, in the final convolutional layer and used global average pooling across the 3 dimensions of the feature map to compute weights for GRAD-RAM, 𝛼_𝑚_:

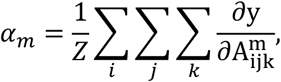

where 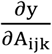 is the gradient of predicted BMI with respect to the 𝑖^𝑡ℎ^, 𝑗^𝑡ℎ^, 𝑘^𝑡ℎ^ unit of the 𝑚^𝑡ℎ^ feature map, where 𝑖 indexes the total height (𝑢), 𝑗 indexes the total width (𝑣), and 𝑘 indexes the total depth (𝑤). Once the weight was computed for each feature map, a weighted combination of the feature maps was used to develop the heatmap, 𝐻 ∈ ℝ^𝑢,𝑣,𝑤^:

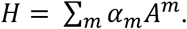

The heatmaps were zoomed to match the original image size using cubic spline interpolation(Virtanen et al., 2020). The heatmap was then normalized to have zero mean and unit variance and all values below 2 were set to zero in order to identify the most important regions to predict, as was done in Vakli et al. (2020). Note that in the original GRAD-CAM, a ReLU function would be applied to the heatmap prior to normalization to identify only the regions of the image with a positive influence on class prediction.

However, in regression tasks, we care about both positive and negative influences on the predicted outcome, and therefore, the ReLU function was not applied to the heatmap for GRAD-RAM (Vakli et al., 2020). The final heatmap was overlayed on the MNI-152 template using open-source software, MRIcroGL (https://www.nitrc.org/projects/mricrogl), and important brain regions were identified via the Automated Anatomical Labeling atlas (Rolls et al., 2020).

We used BMI cut-offs to categorize subjects into the following two groups: healthy BMI/predict healthy BMI (H/pred H) and overweight BMI/predict overweight BMI (O/pred O). The healthy category was defined as having a BMI between 18.5 and 25 and the overweight category was defined as having a BMI greater than or equal to 25. Heatmaps were generated for subjects in the ‘lockbox set’ belonging to each category and averaged across subjects to assess whether the model focused on different brain regions for predictions of healthy BMI compared to predictions of high BMI. By examining the heatmaps for concordant prediction (e.g., obese participants being correctly identified as obese), we can identify regions in the brain associated with accurate BMI classifications for those belonging to the healthy weight and overweight categories. Note that no subjects in the data fit the underweight description of an observed BMI less than 18.5 which is why we did not calculate the heatmaps for those observed to have underweight BMI.

#### 2.5.2 t-Distributed Stochastic Neighbour Embedding

We used t-distributed Stochastic Neighbour Embedding (t-SNE), to determine whether the 3D-CNN was in fact learning discriminative patterns in the brain associated with BMI. T-SNE is a visualization technique for high-dimensional data which assigns observations in the data a location in a 2- or 3- dimensional space while preserving the distance between observations in their original space (Van der Maaten & Hinton, 2008). Therefore, to assess whether the 3D-CNN learned discriminative patterns, we plotted the 255 pre-engineered 3D-CNN features in a 2- dimensional space and assessed whether observations with similar BMI lied closer to one another in space. We repeated this visualization technique for the pre-engineered features derived from the sMRI regions of interest to see how results compared between the two feature sets. The visual outputs of the t-SNEs were examined with different perplexity values and at different random seeds to ensure that the overall conclusions were consistent across the varying parameters.

As a further analysis to explain the AI results, we also investigated whether the DL-CNN neuroanatomical profile of BMI could be attributed to factors such as sex, age, intra-cranial volume, delay discounting, general cognitive ability, depression, anxiety, alcohol use, and motor skills through t-SNE. We assessed whether any patterns emerged in these demographics and cognitive outcomes across the observations in the 2-dimensional space to see whether an association exists between the 255 3D-CNN features and the respective outcome.

## 3 Results

### 3.1 3D-CNN outperforms machine learning models and Tabnet in predicting body mass index

Tables 1 and 2 present the evaluation metrics for the 3D-CNN, random forest, elastic net, and TabNet models across CV folds and on the lockbox set, respectively. The 3D-CNN fit to 3D sMRIs achieved the highest prediction performance among all models evaluated with a mean 𝑅^2^ = 0.325 across the 15 CV folds and 𝑅^2^ = 0.442 on the lockbox. In comparison, the random forest fit to atlas-based ROI variables achieved a mean 𝑅^2^of 0.063 during CV and an 𝑅^2^ of 0.031 on the lockbox set, making it the second-best performing model.

**Table 1.** Mean (SD) prediction accuracy metrics across repeated (x3) 5-Fold cross-validation for random forest, elastic net, TabNet, and 3D-CNN in the prediction of BMI in healthy young adults with best performing model in bold.

| Model | Predictors | MSE | MAE | R2 |
| --- | --- | --- | --- | --- |
| Random forest | Covariates | 25.269 (5.443) | 3.912 (0.441) | 0.021 (0.045) |
|  | Brain | 23.900 (6.252) | 3.849 (0.528) | 0.065 (0.172) |
|  | All | 24.292 (5.718) | 3.897 (0.446) | 0.051 (0.145) |
| Elastic net | Covariates | 25.589 (5.708) | 3.964 (0.463) | 0.011 (0.045) |
|  | Brain | 25.170 (5.172) | 3.972 (0.420) | 0.022 (0.074) |
|  | All | 24.465 (5.494) | 3.889 (0.458) | 0.052 (0.087) |
| TabNet | Covariates | 25.632 (5.956) | 3.894 (0.514) | 0.010 (0.068) |
|  | Brain | 26.130 (4.514) | 4.006 (0.347) | -0.021 (0.072) |
|  | All | 26.704 (5.884) | 3.998 (0.480) | -0.035 (0.101) |
| **3D-CNN** | **sMRI** | **16.713 (4.240)** | **3.119 (0.395)** | **0.334 (0.145)** |

**Table 2.** Prediction accuracy Metrics on ‘lockbox’ dataset for random forest, elastic net, TabNet, and 3D- CNN in the prediction of BMI in healthy young adults with best performing model in bold. Model from best performing CV fold used for prediction.

| Model | Predictors | MSE | MAE | R2 |
| --- | --- | --- | --- | --- |
| Random forest | Covariates | 3.972 | 28.231 | 0.043 |
|  | Brain | 4.037 | 28.583 | 0.031 |
|  | All | 3.956 | 28.318 | 0.040 |
| Elastic net | Covariates | 28.415 | 3.983 | 0.037 |
|  | Brain | 28.663 | 4.062 | 0.029 |
|  | All | 26.929 | 3.961 | 0.087 |
| TabNet | Covariates | 29.635 | 3.899 | -0.005 |
|  | Brain | 28.659 | 3.972 | 0.029 |
|  | All | 26.873 | 3.852 | 0.089 |
| 3D-CNN | sMRI | **16.438** | **2.928** | **0.442** |

Despite the atlas-based ROI machine learning models not performing as well as the 3D-CNN, a brain-BMI association was still apparent. That is, elastic net and random forest both performed better when fit to data comprising both atlas-based ROI features and covariates in comparison to data comprising only covariates. While both the random forest and elastic net achieved an 𝑅^2^ of 0.021 and 0.011, respectively, when fit to covariate-only data, the 𝑅^2^s increased to roughly 0.05 for both models once ROI variables were included. On the other hand, the prediction performance of TabNet decreased with the addition of the atlas-based ROI variables, with the 𝑅^2^dropping from 0.010 to -0.035, compared to the covariate-only model.

Prediction performance of both random forest and elastic net substantially improved when fit to data comprising the 3D-CNN engineered features (see Tables 3 and 4 for CV and lockbox performance, respectively). Specifically, the random forest achieved a mean 𝑅^2^ of 0.383 across CV folds and an 𝑅^2^ of 0.485 on the lockbox set while elastic net achieved a mean 𝑅^2^ of 0.395 across CV folds and a 𝑅^2^ of 0.456 on the lockbox set. Evaluation metrics remained similar when including covariates in the models in addition to the 3D-CNN features.

**Table 3.** Mean (SD) prediction accuracy metrics across repeated (x3) 5-Fold cross-validation for random forest, elastic net, TabNet, and 3D-CNN in the prediction of BMI in healthy young adults with features engineered by 3D-CNN.

| Model | Predictors | MSE | MAE | R2 |
| --- | --- | --- | --- | --- |
| Random forest | Brain | 15.666 (3.105) | 3.033 (0.325) | 0.383 (0.104) |
|  | All | 15.626 (3.050) | 3.037 (0.318) | 0.385 (0.098) |
| Elastic net | Brain | 15.392 (3.130) | 3.022 (0.345) | 0.395 (0.099) |
|  | All | 15.034 (3.114) | 2.978 (0.360) | 0.410 (0.091) |

**Table 4.** Prediction accuracy metrics on lockbox Dataset for random forest and elastic net in the prediction of BMI in healthy young adults with features engineered by 3D-CNN.

| Model | Predictors | MSE | MAE | R2 |
| --- | --- | --- | --- | --- |
| Random forest | Brain | 15.182 | 2.862 | 0.485 |
|  | All | 15.217 | 2.867 | 0.484 |
| Elastic net | Brain | 16.048 | 2.959 | 0.456 |
|  | All | 16.053 | 2.960 | 0.456 |

#### 3.2 Explainable AI identifies and explicates key regions in 3D-CNN prediction

### 3.2.1 Localization Heatmaps via GRAD-RAM

The mean localization maps produced by GRAD-RAM for those in the H/pred H category and the O/pred O category are presented in Figure 3. Overall, brain regions with greater influence on predicted BMI were similar for the two groups. In both groups, the brain regions relevant to the trained 3D-CNN included part of the occipital lobe (calcarine sulcus, lingual gyrus), the cerebellum (lobules 3, 4_5, 6, vermis 3, vermis 4_5, vermis 6), and the basal ganglia (caudate nucleus, putamen, globus pallidus) in addition to the fusiform gyrus, the transverse temporal gyrus, the thalamus, precuneus, the posterior cingulum and the hippocampus. Brain regions highlighted by the heatmap were predominately in the right hemisphere with the exception of the posterior cingulum. The insula was associated with healthy BMI but not with overweight BMI, whereas the inferior temporal gyrus and parahippocampal gyrus were implicated in overweight BMI but not in healthy BMI. Additionally, specific subregions varied between the groups. For instance, the cuneus, part of the occipital lobe, was highlighted only in heatmaps of healthy individuals. In contrast, the heatmap for overweight individuals encompassed more regions of the cerebellum, such as cerebellum lobules 8, 9, 10, and crus 1, compared to healthy individuals.

**Figure 3.**
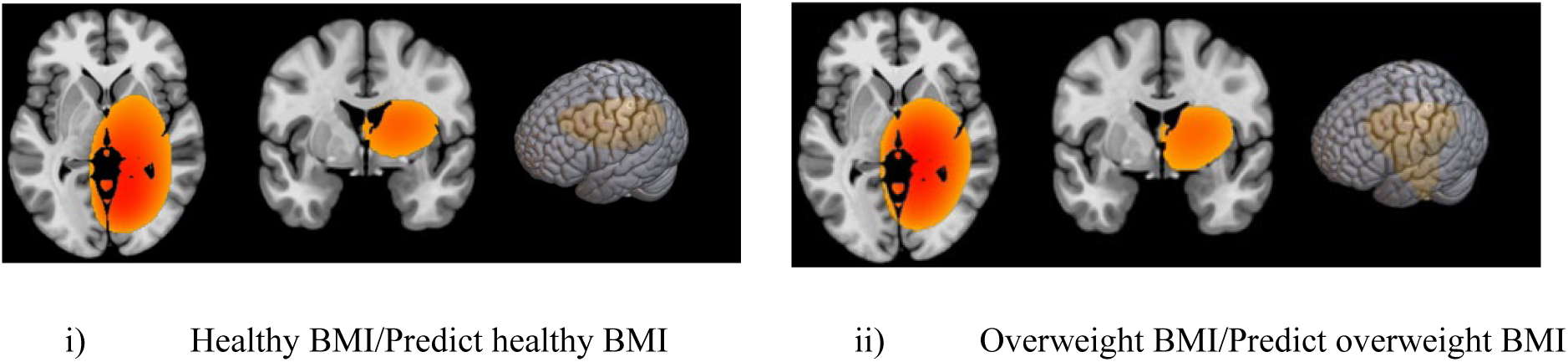
Heatmaps Produced by GRAD-RAM Identifying Relevant Brain Regions in the Prediction of BMI for Categories i) H/Pred H (18.5 < BMI < 25) and ii) O/pred O (BMI ≥ 25). Heatmaps Overlayed on the MNI-152 Space with Darker Red Indicating Greater Importance to BMI Prediction and Yellow Indicating Less Importance. Primary regions in heatmap include i) part of the occipital lobe, the cerebellum, and the basal ganglia and the fusiform gyrus, thalamus, precuneus, insula, and the hippocampus in the right hemisphere; and ii) part of the occipital lobe, the cerebellum, and the basal ganglia and the fusiform gyrus, thalamus, inferior temporal gyrus, precuneus, hippocampus, and parahippocampal gyrus in the right hemisphere.

### 3.2.2 3D-CNN Feature Visualization via t-SNE

Figure 4 presents the visualization of BMI across observations in the 2-dimensional space based on i) the 255 3D-CNN features and ii) the 153 atlas-based ROI features. A clearer pattern of BMI emerged across observations in the 2-dimensional space based on the 3D-CNN features compared to the 2-dimensional space based on the atlas-based ROI features confirming that the 3D-CNN features from the final convolutional layer were more effectively able to discriminate patterns in the brain associated with BMI. More specifically, in the 2D space based on 3D- CNN features, those with higher observed BMI tended to have lower scores on the first dimension than those with lower BMI. In contrast, in the 2D space based on the handcrafted ROI features, no patterns emerge regarding BMI.

**Figure 4.**
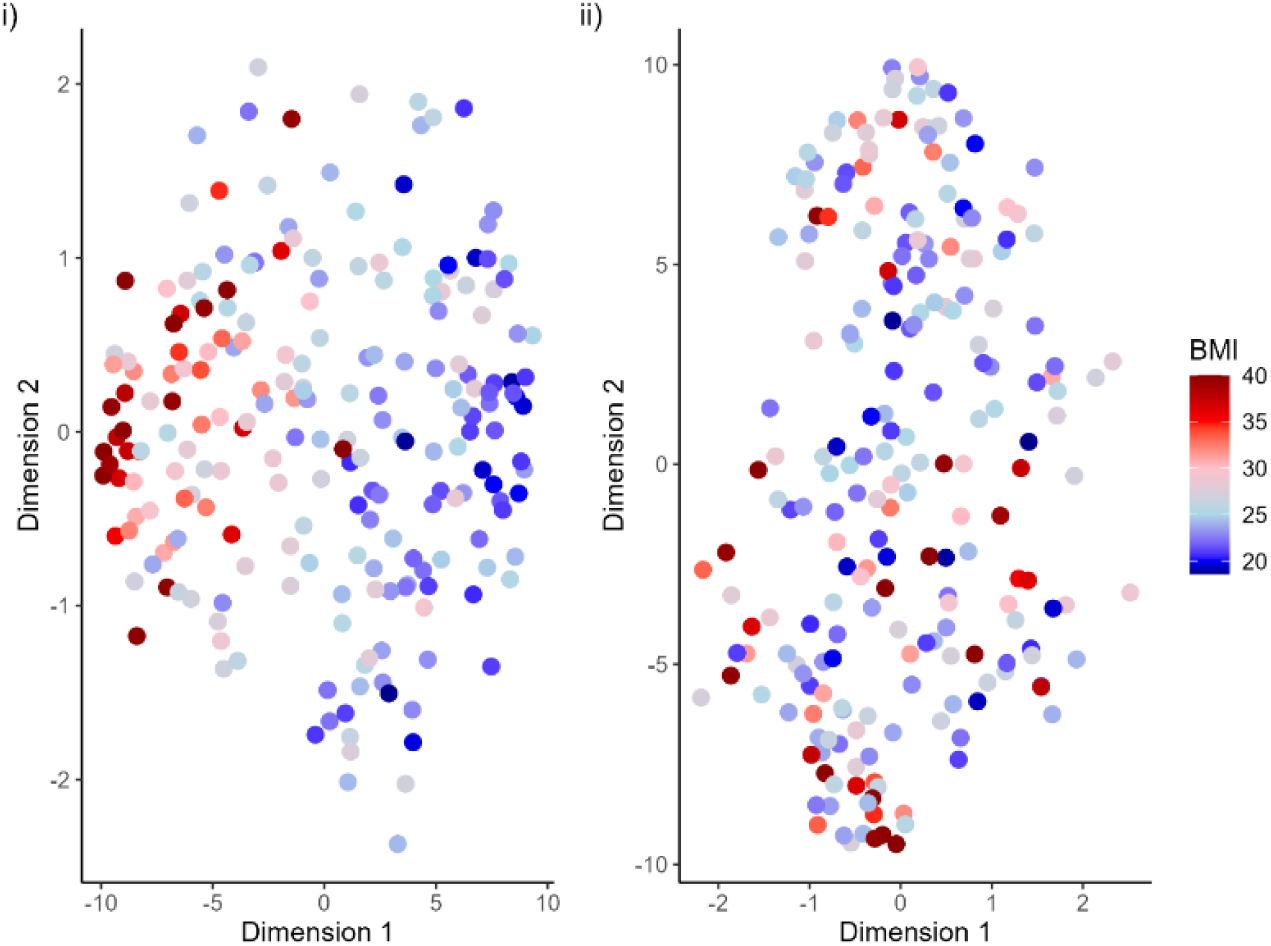
Patterns of Observed BMI across Projections of i) 255 3D-CNN Features and ii) 153 Atlas-based ROI features in a 2D Space via t-SNE.

There were moderate associations between the 2D scores based on the 3D-CNN features and high reward magnitude delay discounting, general cognitive ability, motor skills (gait speed, dexterity), and frequency of alcohol use in the past year. In particular, those with more impulsive delay discounting, lower general cognition ability, greater alcohol use, lower gait speed and worse dexterity tended to have lower scores on dimension 1 (see Figure S1 in Supplemental). These moderate associations are not surprising given the small, negative correlation between observed BMI and delay discounting (Spearman’s 𝜌 = −0.252, 𝑝 < 0.001), total cognition score (𝑟 = −0.269, 𝑝 < 0.001), gait speed (𝑟 = −0.143, 𝑝 = 0.035), and dexterity (𝑟 = −0.233, 𝑝 < 0.001). However, interestingly, observed BMI only had a small, non-significant association with frequency of alcohol use (Spearman’s 𝜌 = 0.097, 𝑝 = 0.167). No significant associations were present across observations in the 2D space based on 3D**-**CNN features for age, sex, anxiety, depression, alcohol dependence, or low reward magnitude delay discounting (see Figures S2-S8 in Supplemental).

## 4 Discussion

Our study contributes to the growing body of literature illuminating the relationship between the brain and obesity (Adise et al., 2021; Guan et al., 2020; Vakli et al., 2020; Wang et al., 2023; Xu, Owens, et al., 2023). The results demonstrate that the 3D-CNN massively outperformed other forms of machine learning and deep learning methods that rely on tabular atlas-based ROI features in the prediction of BMI. The ability of the 3D-CNN to learn representations in the sMRIs is likely to have contributed to this improved performance, suggesting that BMI representations in the brain are too complex and abstract to be explained by atlas-based ROI features. Further evidence for this comes from the sequential boosting approach where we demonstrated substantial improvement of machine learning methods, elastic net and random forest, when fit to data comprising the 3D-CNN engineered features thereby providing evidence that the original difference in performance between the models came down to the features used, not the models themselves. It is important to note that atlas-based ROI features are subject to the *brain concordance problem*, meaning that the parcellation and region names are sensitive to the chosen atlas (Bohland et al., 2009), complicating research on the brain-BMI connection. Allowing deep learning models to learn representations of the brain directly from 3D images is a potential solution to this problem when looking for associations between brain structure and a given outcome.

While earlier studies provided preliminary evidence of the utility of 3D-CNN in the prediction of BMI without cross-validation or a lockbox dataset, we were able to show that the brain-BMI association is robust enough for high performance in a relatively modest sample (*n* < 1,000) using gold-standard cross-validation and an independent lockbox. Indeed, prediction accuracy was notably higher in the lockbox sample. Overall, acknowledging that the data are cross-sectional and causality cannot be inferred, our results provide convincing evidence of a robust associations between brain morphometry and obesity.

Similar to the findings of Xu et al. (Xu, Owens, et al., 2023), we did not find evidence to suggest that the association between the brain structure and BMI is confounded with demographics. First, the machine learning models fit to data comprising the 3D-CNN features substantially outperformed the models fit to data comprising covariates only. Second, the machine learning models fit to data comprising the 3D-CNN features in addition to covariate data performed similarly to the models when fit to data comprising only the 3D-CNN features. Therefore, it does not appear that superficial characteristics can explain the relationship between the 3D-CNN features and BMI. Moreover, the t-SNE visualizations did not show any patterns of age, sex, or intra-cranial volume across observations in the 2D space based on the 3D-CNN features.

Explainable AI methods revealed the regions in the brain relevant to the neuroanatomical profile of BMI. Interestingly, the 3D-CNN focused on regions of the brain that play a role in cognition, reward processing, motor control, emotional processing, and sensory processing. The localization heatmaps identified the caudate, putamen, and pallidum as potentially relevant brain regions to the prediction of BMI. These three brain regions are part of the basal ganglia, a dopaminergic system which plays an important role in motor control and has been implicated in conditions associated with overconsumption, such as drug addiction (Morais-Silva & Lobo, 2024; Salokangas et al., 2000; Xu, Xu, et al., 2023) and obesity (Takeuchi et al., 2020; Tan et al., 2022). In addition, the heatmaps suggested that the hippocampus and thalamus were both relevant to BMI prediction which coincides with the finding that these make up a network along with the hypothalamus that is responsible for energy metabolism and weight management (Davidson et al., 2005; Davidson & Jarrard, 1993). The 3D-CNN could have dissected information relevant to this network that may not be available when only considering the atlas-based ROI features.

Another interesting finding is that the regions identified as being important to the 3D-CNN prediction included the hippocampus and the parahippocampal gyrus, which have been previously linked to delay discounting (Frost & McNaughton, 2017). Given that the relationship between delay discounting and BMI is well established (Amlung et al., 2016; X. Liu et al., 2022; Y. Liu et al., 2019; Weller et al., 2008), it is not surprising that the structural influences in the brain might be similar between the two outcomes. Indeed, for the same reason, these findings suggest potential neurobiological markers of obesity as candidate treatment targets for repetitive transcranial magnetic stimulation, which has been shown to be effective for treating substance use disorder (Harel et al., 2022; Zangen et al., 2021), another condition of overconsumption, potentially by way of delay discounting (Shevorykin et al., 2022). However, it is important that these results are interpreted with caution given that the spatial resolution toward the end of the network is poor after multiple layers of max pooling. While GRAD-RAM is an effective method to identify clusters relevant to prediction, the loss of finer details in the image may lead to imprecision in identifying brain regions.

In terms of considerations, noteworthy strengths of the study include: 1) thorough investigation of whether 3D-CNN improves prediction of BMI compared to machine learning models; 2) the assessment of generalizability of results across different train/test splits via nested cross-validation; and 3) evaluation of a fully independent lockbox dataset; 4) explainable AI analyses to reduce the opacity of the 3D-CNN results. On the other hand, limitations include a relatively small sample size for deep learning analyses and a cross-sectional design that prevents conclusions about whether these brain features are etiological causes of obesity or consequences from developing obesity (or recursive relationships in both causal and consequential roles over time). In a longitudinal sample, it would be interesting to see whether misclassifications of BMI category could be predictive of future health outcomes such as obesity and eating disorders similar to how misclassification of brain age is predictive of outcomes such as dementia (Biondo et al., 2022) and Alzheimer’s (Poloni et al., 2022).

In sum, our findings demonstrate the remarkable promise of deep learning models for learning data representations to characterize the relationship between brain morphometry and BMI. Despite the additional complexity of these models, there is substantial improvement in prediction performance from learning features directly from the data rather than using atlas-based ROI features. While the atlas-based ROI features may be useful for simpler outcomes with a clearer connection with the brain, they do not appear to aptly capture all aspects of the brain structure implicated in BMI. Priorities for next steps include extending these findings to different samples, as the results may not generalize to different age groups or demographics, and using deep learning in prospective designs to longitudinally illuminate the role of the brain in the development of obesity.

## Data and Code Availability

sMRI data and the covariates used in this study can be requested from the Human Connectome Project: https://www.humanconnectome.org/. All code for the current study is available at https://github.com/PBCAR/3DCNN_BMI.git.

## Author Contributions

AC, ME, MO, and JM conceived of the presented idea. MO prepared the data and identified the initial analytic strategy. AC determined the final analytic strategy and conducted the analyses; ME assisted with follow-up analyses explaining the results in terms of the brain regions implicated and behavioral processes. AC wrote the original manuscript; MO, ME and JM provided editing and critical review of all content. JM acquired funding and provided supervision.

