## Supplemental Materials for "Deep learning using structural MRI massively improves prediction accuracy of body mass index"

**Figure S1.**

*Patterns of i) Delay Discounting AUC at $40K, ii) Cognition Function Composite Score, iii) Gait Speed, iv) Dexterity, and v) Frequency of Alcohol Use across Projections of 255 3D-CNN Features in a 2D Space via t-SNE.*

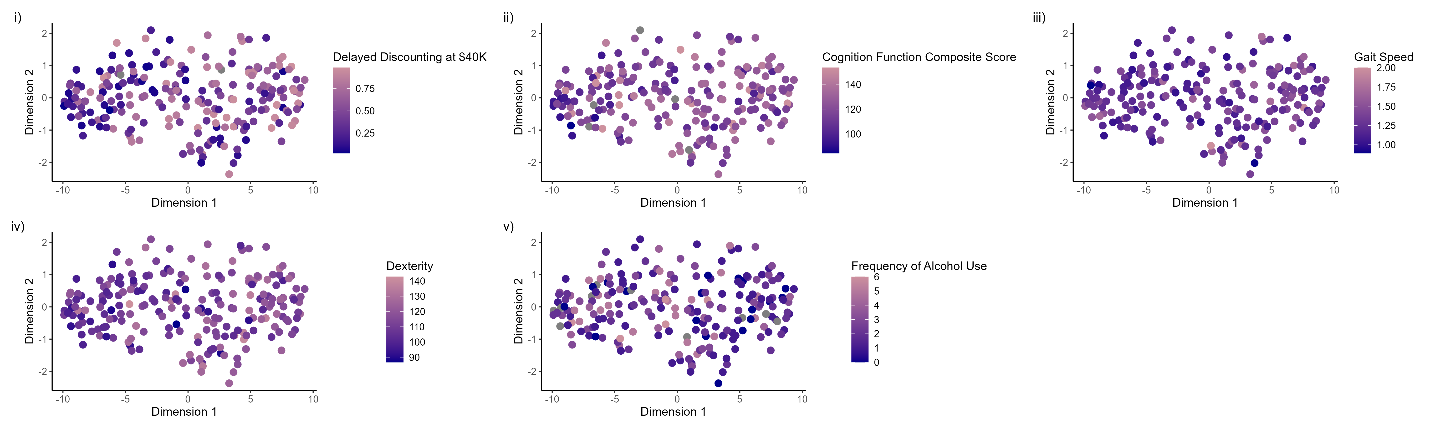

**Figure S2.**

*Age across Projections of 255 3D-CNN Features in a 2D Space via t-SNE.*

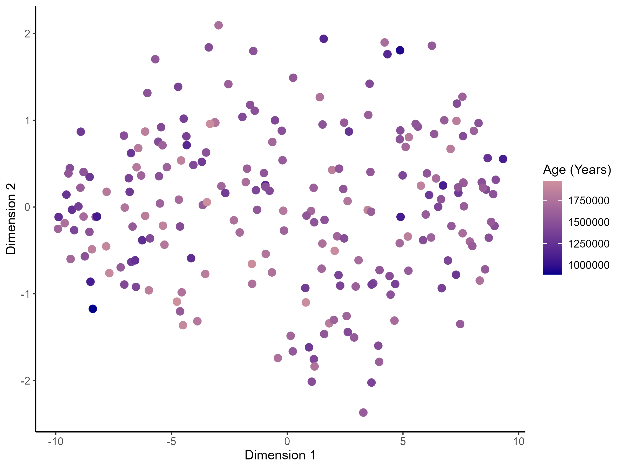

**Figure S3.**

*Intra-Cranial Volume across Projections of 255 3D-CNN Features in a 2D Space via t-SNE.*

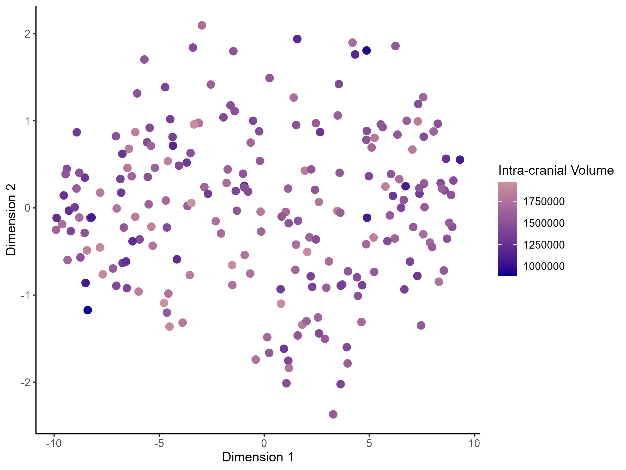

**Figure S4.**

*Sex across Projections of 255 3D-CNN Features in a 2D Space via t-SNE.*

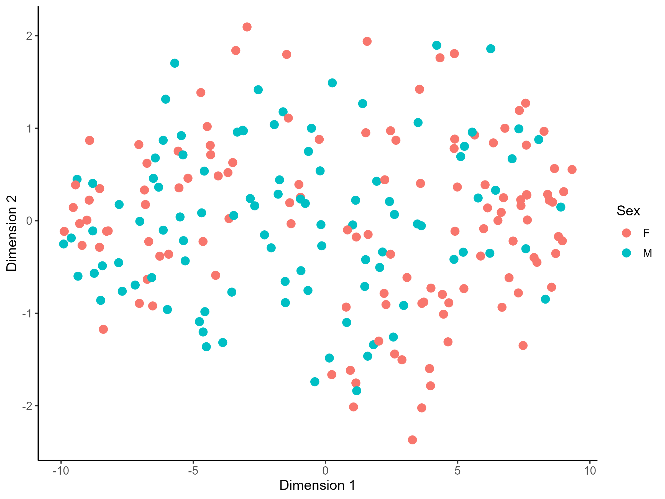

**Figure S5.**

*DSM Anxiety Problems Score across Projections of 255 3D-CNN Features in a 2D Space via t-SNE.*

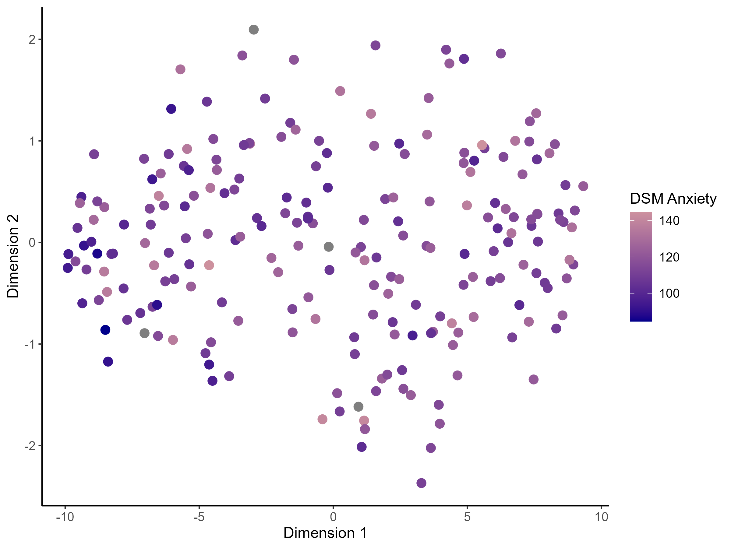

**Figure S6.**

*DSM Depressive Problems Score across Projections of 255 3D-CNN Features in a 2D Space via t-SNE.*

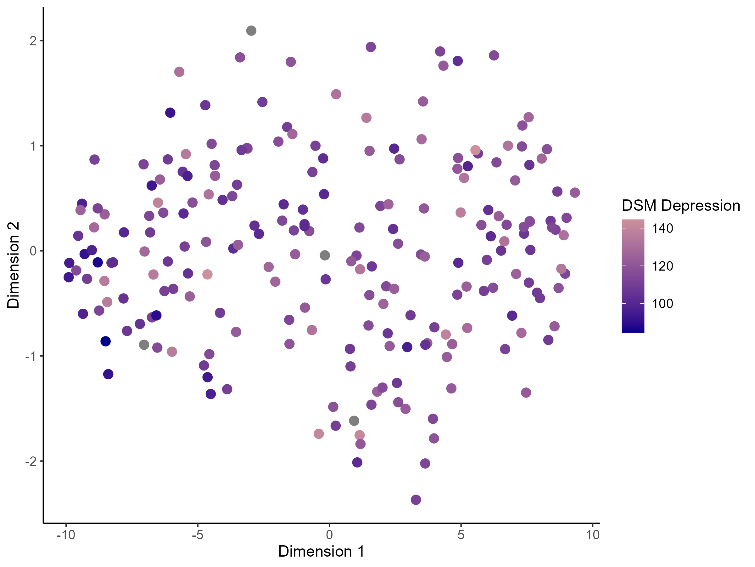

**Figure S7.**

*Delay Discounting AUC at $200 across Projections of 255 3D-CNN Features in a 2D Space via t-SNE.*

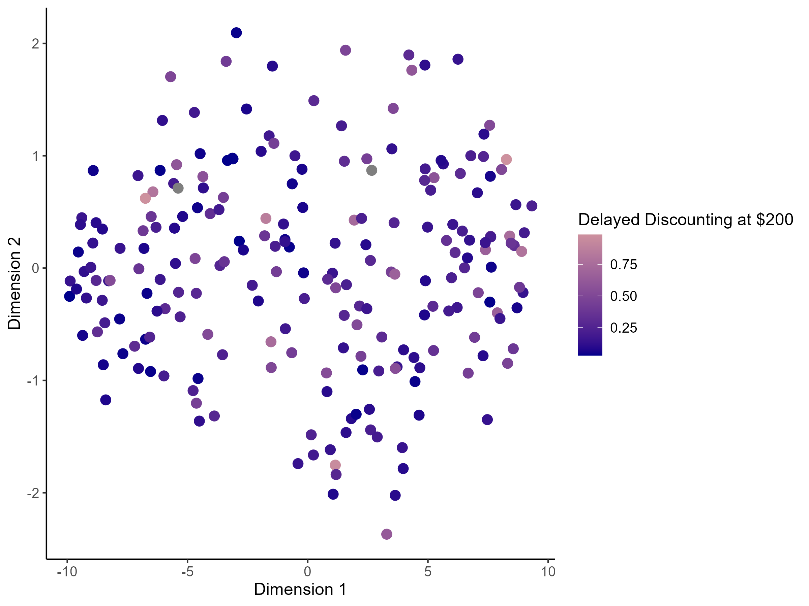

**Figure S8.**

*Alcohol Dependence across Projections of 255 3D-CNN Features in a 2D Space via t-SNE.*

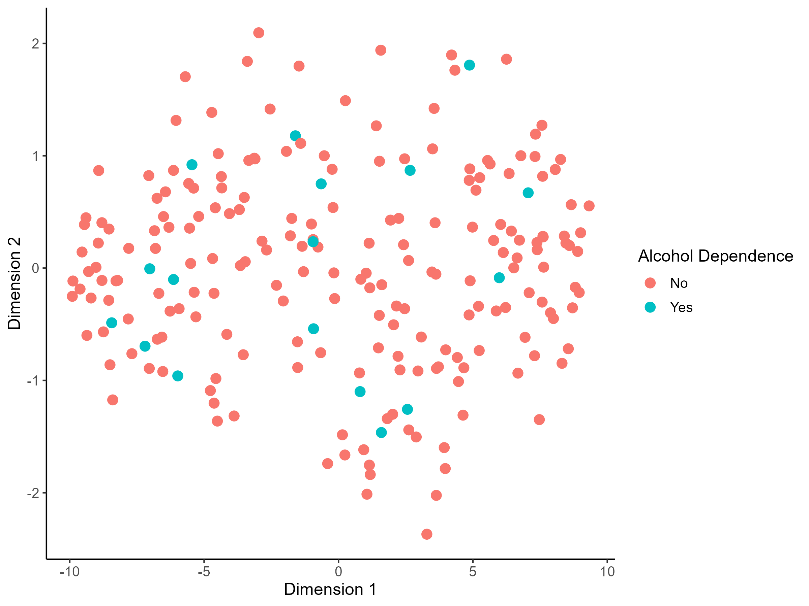
